# On the delay in propagation of action potentials

**DOI:** 10.1101/763698

**Authors:** J. Xu, S. Xu, F. Wang, S. Xu

## Abstract

The signal delay during the propagation of action potentials is one of the key issues in understanding the mechanisms of generation and propagation of neural signals. Here we reanalyzed related experimental data to demonstrate that action potentials in the propagation process along a myelinated axon are highly overlapped in the time scale. The shift in time of two successive signals from neighboring nodes, defined as delay time ***τ*** in this work, is only tens of microseconds (16.3-87.0 μs), thus is only ~ 0.8-4.4 % of the measured average duration of an action potential, ~ 2 ms. This fact may reveal a huge gap to the commonly accepted picture for propagation of neural signal. We could apply the electromagnetic soliton-like model to well explain this phenomenon, and attribute ***τ*** to the waiting time that one signal source (i.e., ion channel cluster at one node) needs to take when it generates an electromagnetic neural pulse with increasing intensity until the intensity is higher than a certain point so as to activate neighboring signal source. This viewpoint may shed some light on a better understanding of the exact physical mechanism of neural signal communication in a variety of biosystems.

**Statement of Significance:** The delay time during the propagation of action potentials is an important term in understanding the mechanisms of generation and propagation of neural signals. In this article we analyzed published experimental data and showed that action potentials from two neighboring Ranvier nodes are highly overlapped in time, with an average shift of tens of microseconds, which occupied only ~ 0.8-4.4 % of the average duration of an action potential (2 ms). The electromagnetic soliton-model seemed the best model to explain this phenomenon.

The viewpoint of this article may shed some light on a better understanding of the exact physical mechanism of neural signal communication, and be tractive to researchers in a variety of fields, such as neuroscience, brain-computer interface, etc..

## Introduction

The nature of signal delay in propagation of action potentials (APs) along an axon is one of the key issues in understanding of the mechanism of generation and transmission of neural signals, but with less attention. The total delay in time during the propagation process of an action potential may be divided into two parts: the delay on transmission path, and that on signal sources, *i.e*., ion channels. There are different perspectives related to this issue. *Saltation with respect to time* (1, 2) attributes the delay in neural signal transmission to the signal relay process at Ranvier nodes, because it claims that the current transmits between nodes with the nearly speed of light. *Saltation with respect to space* points out that the time required for neural signals to propagate on the internode segment is the main part determining the total conduction time (3). With the development of advanced patch clamp technique, it showed that the signal delay at each node was within a narrow range and independent of the internode length (4). This strongly indicated that in propagation of neural signals, the time is spent mostly at Ranvier nodes rather than at internode segments.

In this paper we reanalyzed published experimental data, and figured out a clear picture on the underlying mechanism of the measured propagation speed of action potentials along axons.

## Results & Discussion

### Signal delay in myelinated axons

The propagation speed *v* of action potentials is calculated as

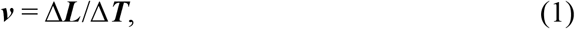

where Δ***L*** is the distance between two points under test and Δ***T*** is the total delay time between these two testing points. The Δ***L*** can be rewritten as

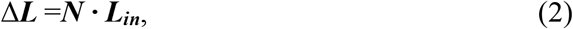

where ***L_in_*** is the average internode length in a myelinated axon. and ***N*** is the number of Ranvier nodes between two testing points. Similarly, Δ***T*** can be rewritten as

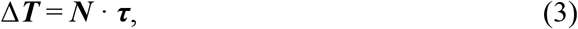

where ***τ*** is the average delay time cost at each node. Thus ***τ*** can be deduced as

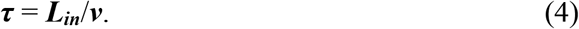

In **Table 1**, we listed some experimental data and our calculated average delay ***τ*** following the above equation (4). The calculated *τ* ranges from 16.3 to 87.0 μs. This is compliant with other experimental results, where a so-called *latency time* of 10-50 μs was measured (4, 5). The delay was also defined as *elapsed time*, and Akaishi (6) reported a calculated value of around 1 μs. In some other literatures it was also named as *internodal conduction time* with a value of 74 μs (7).

**Table 1.**
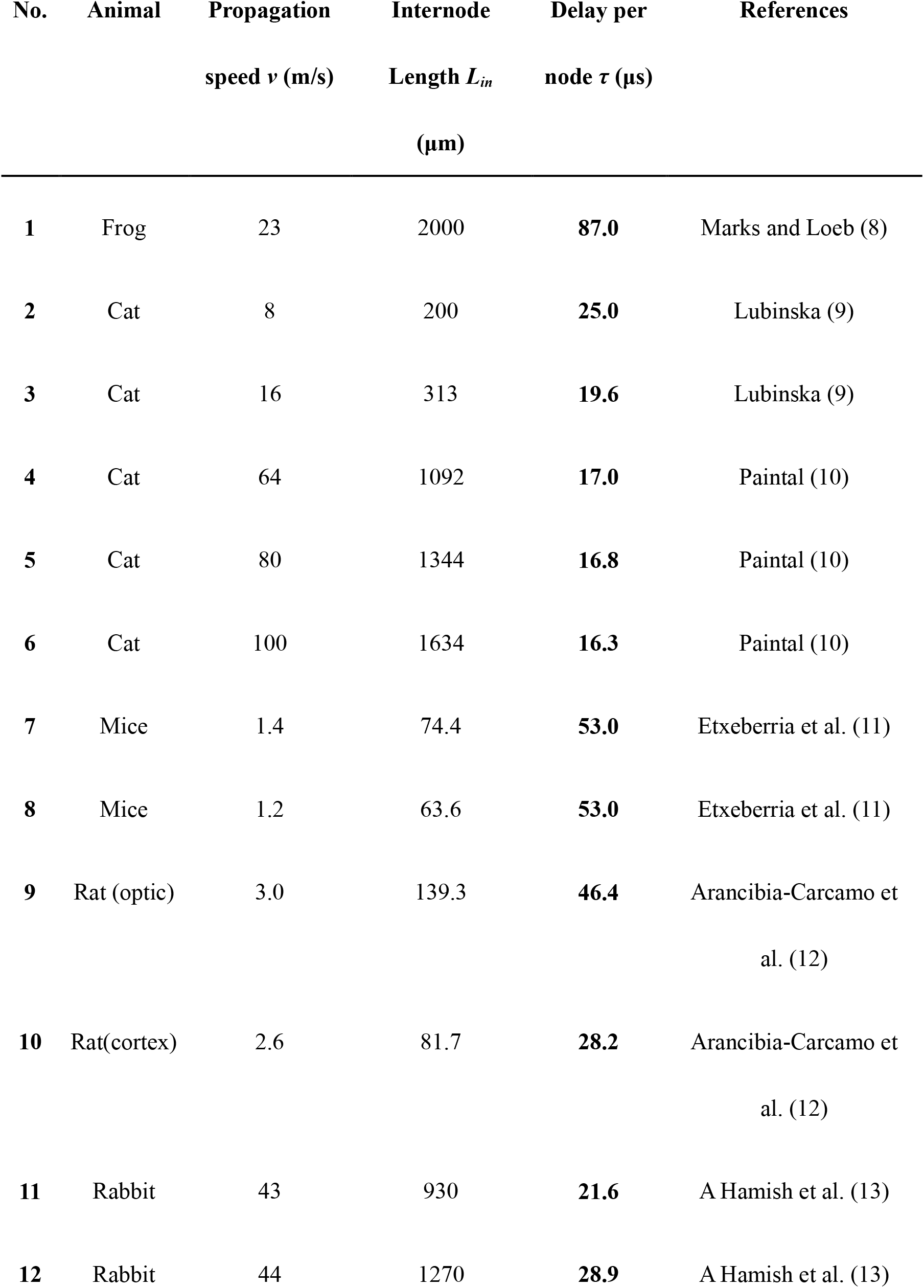
Average signal delay τ

### Illustration of the action potential peaks

Now we have learnt two experimental facts. One is that the action potentials recorded at different nodes are almost identical along the same axon, with a duration of around 2 ms (14). The second fact is that the delay per node is less than 100 μs, as shown in **Table 1. Figure 1** illustrates these two facts together schematically. When the action potentials are measured at different nodes (highlighted with different color), they are highly overlapped. The right enlarged box gives a close look at two peaks from two neighboring nodes, where the shift of the blue curve to the previous green curve in the time, *i.e*., a delay ***τ***, is less than 100 μs, much shorter than the period ***t_0_*** of the action potential (≈ 2 ms).

**Figure 1.**
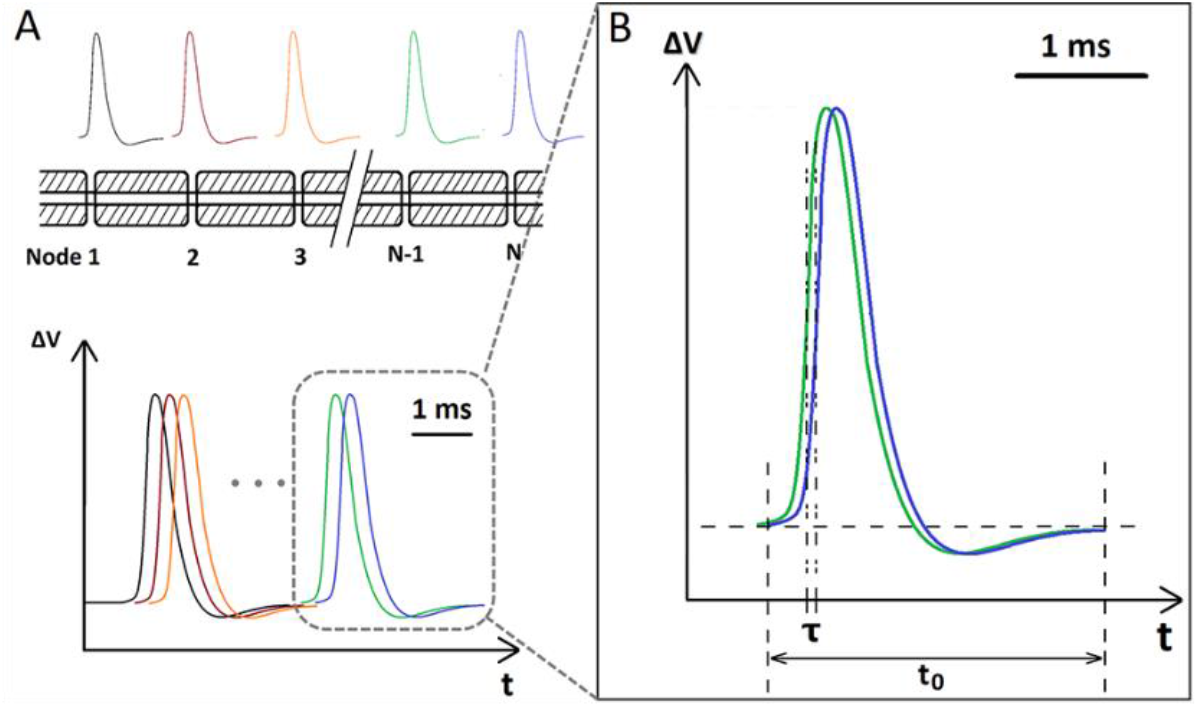
T The illustration for propagation image of action potentials in myelinated axons. A) the schematic diagram of myelinated fibers and their action potentials in the same timeline. B) the shift of two neighboring action potentials in time.

### Origin of signal delay at Ranvier nodes

Here are three main kinds of models for propagation of neural signals. The *electromechanical wave model* where neural signal is mechanical dense waves (15) suggests that the delay of signals mainly costs at internode segments. This contradicts the later experimental results (4, 5). The *cable model* claims the neural electrical signals propagate via the local transverse current field (1). But it seems not valid for myelinated axons. The distance between two neighboring signal sources, *i.e*., Ranvier nodes, is as long as several millimeters, thus much longer than the Debye length (about several nanometers) of ions in an electrolyte solution resulting from screen effect (16). The *electromagnetic soliton-like model* states that the neural signals are electromagnetic pulses generated by the transmembrane transient ion current, and they propagate via the naturally formed softmaterial dielectric waveguides with the structure of “intracellular fluid – lipid membrane – extracellular fluid” (17–19). Both experiments and simulations showed the electromagnetic pulse signal has limited transmission efficiency when propagating via the softmaterial waveguide pathway.

This results in an attenuated field strength when the pulsed signal arrives at the next signal source (20, 17, 19, 21). This model explains well the highly overlapped successive action potentials as shown in **Figure 1**.

**Figure 2** schematically illustrates electromagnetic soliton-like model for the origin of signal delay *τ* between two neighboring nodes, namely *Node 1* and *Node 2*. A typical Δ***V-t*** curve measured at *Node 1* is plotted in **Figure 2A**. It is well known that there exists a threshold point ***V_threshold_*** at ***t***=***t_0_***, under which, the local ion channels at *Node 1* are closed. At this time, the ion channels at *Node 2* are still closed, although they have sensed an attenuated electric filed from *Node 1*. **Figure 2B** illustrates an image of this attenuated signal at *Node 2*. As the transmembrane ion current increases its strength at *Node 1* thus resulting in an abrupt increment of local electric field, the signal amplitude sensed at *Node 2* also increases rapidly. When it reaches the threshold, the ion channels at *Node 2* are triggered open (***t***=**t_2_**) and thus generating its own local pulsed signal, as shown in **Figure 2C**. Note that the curve in **Figure 2B** starts from time **t_0_**, while the curve in **Figure 2C** starts from time **t_2_**. Therefore, the whole Δ***V-t*** curve in **Figure 2D** recorded as action potential at *Node 2* is an overlapping of two curves: one from the attenuated signal transmitted from *Node 1* and the other generated by the local ion channels, which is almost the same as that at *Node 1*. In short, the difference, **t_2_ – t_1_**, is the delay time ***τ*** between these two signals, where **t_1_** and **t_2_** are the open time for *Node 1* and *Node 2* respectively.

**Figure 2.**
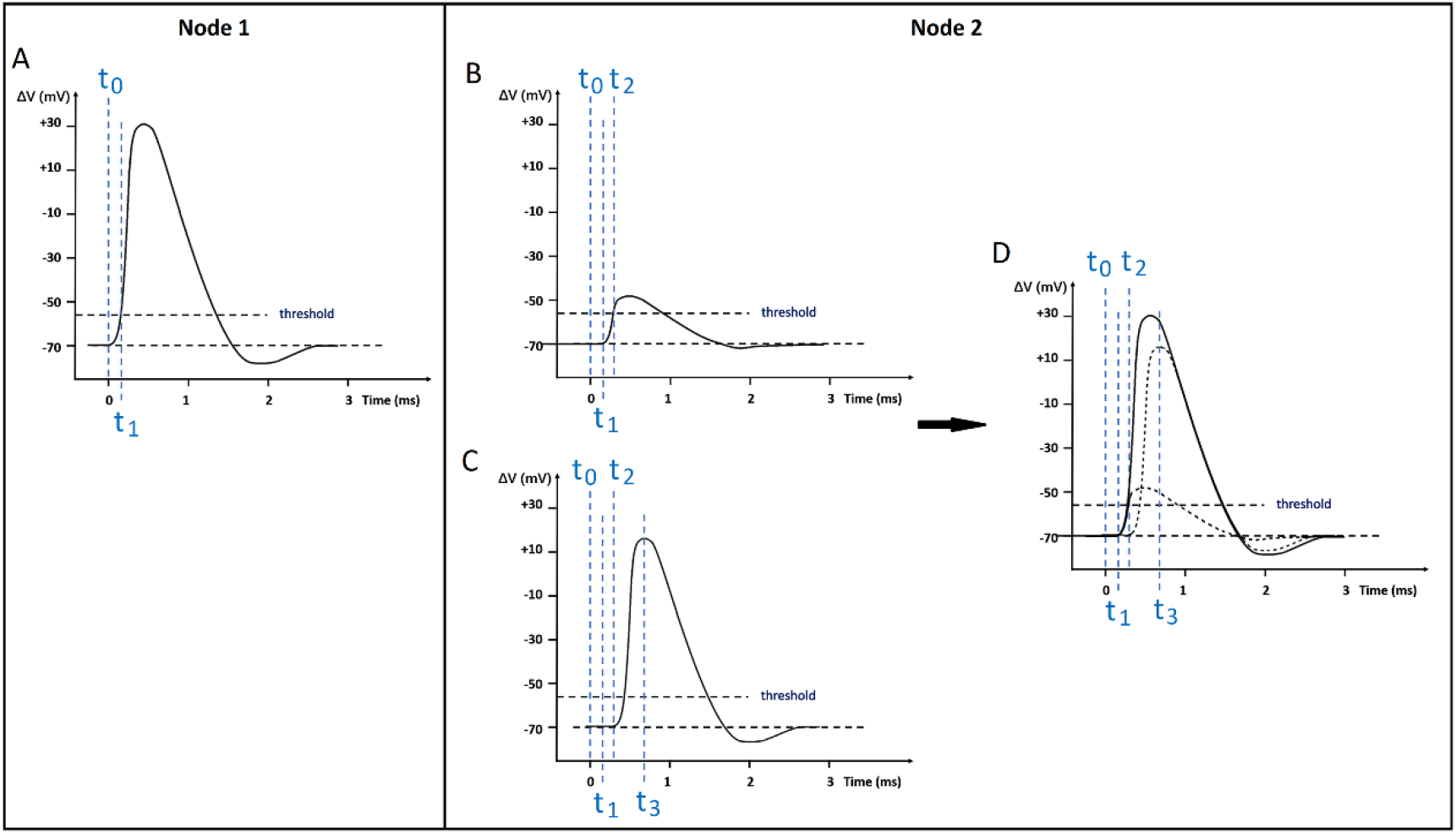
Schematic illustration for the origin of delay time per node. It has two parts: the small one is the attenuated pulsed signal (B) generated by ion channels at *Node 1* (A), and the other (C) is generated by local ion channels at *Node 2* once the attenuated pulse goes up to the threshold to activate them.

**Figure 2** also gives the difference among three terms in time: the delay time per node, the open time of ion channels and the duration of action potential, which is necessary and helpful to understand the process of neural signals’ propagation. The delay time per node, ***τ***, includes two parts. One is the transmission time of neural pulsed signals along internode segment, which can be overlooked due to the near-light speed of electromagnetic pulses. The other is the waiting time that the signal generated by *Node 1* takes to rise from zero to the specific value whose attenuated amplitude reaches to the activation threshold at *Node 2*. Note that the calculated range of ***τ***, 16.3-87.0 μs, is close to the reported activation time constant of voltage-gated sodium channels, 64 ± 6 μs at 100kHz (22). This indicates the inevitable correlation between the delay process and the activation of ion channels. The open time of ion channels, *i.e*., ***t_3_-t_2_*** in **Figure 2**, refers to the lasting time that the channels open, with a range of 0.4-1.0 ms (14). Action potentials include the under-threshold period, depolarization period (activation of sodium channels and the following inwards ion flux), repolarization period (inactivation of sodium channels and activation of potassium channels) and recovery period (return to normal permeability under the working of sodium-potassium ATPase pumps). Therefore, the action potential is actually the apparent image resulting from the inward and outward flows of an amount of ions, rather than the physical essence of the neural signal (6).

This model indicates that the propagation speed of neural signals increases with the internode length ***L_in_***, consistent with experimental discovery (13, 23). For the certain length of an axon, a longer ***L_in_*** results in smaller number of nodes, thus a small value of the total delay time. One possible way to enlarge ***L_in_*** is to increase the thickness of myelin sheath. Both simulation and experiments on large scale softmaterial waveguides showed that, in general the thicker the dielectric layer, the lower the attenuation and thus longer effective transmission length (17), which probably decides the internode length. This is consistent with observations in biosystems (24). Another feasible way to enlarge ***L_in_*** is to enhance the total intensity of signals at one node by increasing the density of ion channels. Indeed, a ion density of 1,000/μm^2^ was observed (25), higher than that in unmyelinated axons, typically 50/μm^2^ (22).

### Is this model valid in neural signal transmission along unmyelinated axons?

In the evolution of living organisms, myelinated axons are developed from unmyelinated axons. Therefore, it is reasonable that the propagation of neural signals in both unmyelinated and myelinated fibers share some common mechanisms. We may apply the scenario of signal delay in myelinated axons to the propagation of neural signals in unmyelinated axons. There may be three major kinds of distribution patterns of voltage-gated ion channels on an unmyelinated axon, as shown in **Figure 3**. The first is a uniform distribution where the ion channels are separated with an average spacing, as reported by Black et al. (26). This spacing should be as long as possible so as to increase the propagation speed of neural signals. Meanwhile, it should not be too long, in order that after attenuation the signal received at neighboring channels could reach the triggering threshold. Obviously, when the original signal strength is increased, it favors a longer spacing between two signal sources. As a result, making a cluster of ion channels as the signal source is probably a better distribution form for ion channels, such as point clusters and ring-like clusters. There are more and more evidences clarified the cluster distribution of ion channels (27, 28). For example, Jahnsen and Llinás (29) found that Na+ ion channels are localized in clusters and the distance between them is about 5-15 μm in aplysia axonal membrane.

**Figure 3.**
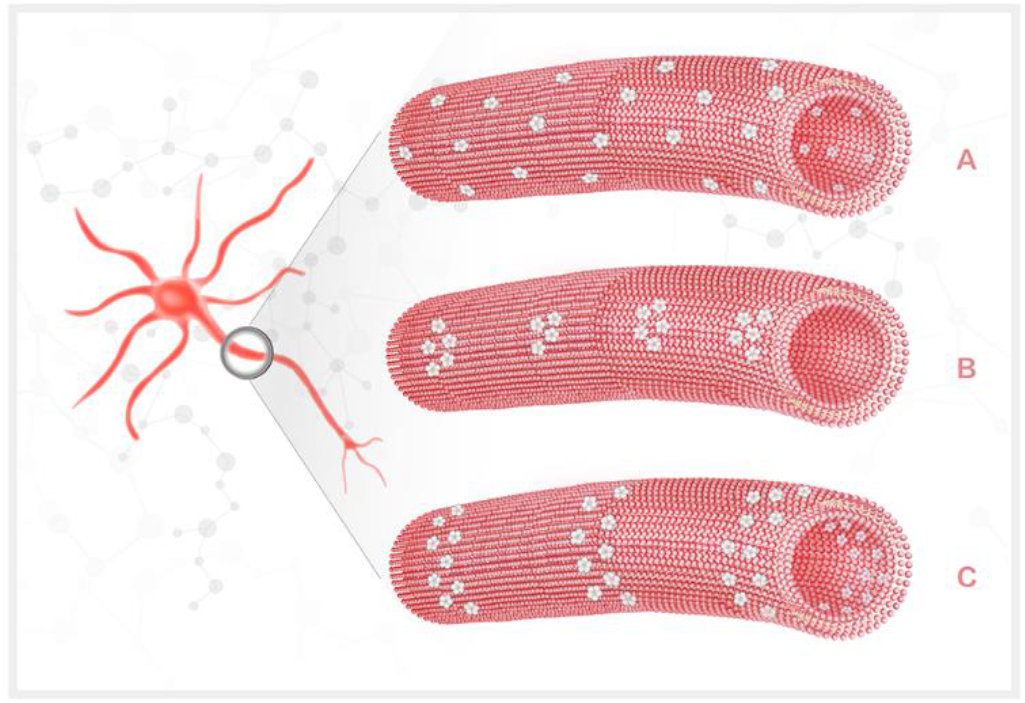
Three possible distribution of voltage-gated ion channels on unmyelinated axons: uniform distribution (A), point cluster pattern (B) and ring-like pattern (C).

Considering the typical propagation speed in unmyelinated axons (1 m/s) and taking 10 μs as the delay time, then there are 100,000 sites of signal source along one-meter long axon. The distance between two adjacent signals sources, l, is then 10 μm, consistent with the reported 5-10 μm (30). Due to uniform attenuation of neural electromagnetic signals in unmyelinated axons where the dielectric layer is the single bilayer lipid, the longer ***l*** could only result from the number, ***M***, of ion channels at one signal source, which can lead to strong enough electromagnetic signals to trigger the neighboring channels. One feasible way to improve ***M*** is to enlarge the diameter of axons. Previous experiments demonstrated that there are always more ion channels on thicker axons(31). Some other studies also reported that the thicker axon usually has larger cluster area (32), more sodium ion channels at signal sources, and thus stronger neural signals.

## Conclusion

In summary, we reanalyzed a variety of experimental data on the propagation of action potentials in myelinated axons, and discussed several different models reported in the literatures. We showed that action potentials generated at neighboring Ranvier nodes, though clearly separated in space, are indeed highly overlapped in time. The shift in time of any two successive signals is less than 100 μs, thus is much less than 10% of the average period of an action potential (~ 2 ms). This shift in time, defined as delay time *τ* in this work, is attributed to the waiting time for the signal source at one node (i.e., ion channel clusters) to sense an attenuated electromagnetic field up to the threshold level. The attenuated electromagnetic fields are generated from the previous node and they transmit through the internode in a speed close to light speed, as according to the electromagnetic soliton-like model. The novel view for the propagation of neural signals may also be applied to unmyelinated axons, and help to better understand the electrical signals communication in neural systems and brains.

## Author Contributions

Jingjing Xu made a significant contribution to the analysis and interpretation of data and phenomenon, and drafted the article. Sanjin Xu and Fan Wang mainly participated in the review of lots of related literatures. Shengyong Xu gave the original concept of the article and revised it. All authors approved the final version of the manuscript.

## Acknowledgement

This work was financially supported by the National Key R&D Program of China (No. 2017YFA0701302) and the research fund from Shandong University (2018GN030). The authors declare that no competing interests exist.

